# Daylight exposure improves sleep and subsequent sleepiness

**DOI:** 10.64898/2026.10.22.761062

**Authors:** Minqi Yang, Thomas van Bergeyk, Roelof A. Hut, Renske Lok

**Affiliations:** School of Education and Cognitive Science, Zhengzhou University, Zhengzhou, 450001, China; Chronobiology Unit, Neurobiology Expertise Group, Groningen Institute for Evolutionary Life Sciences, University of Groningen, Groningen, Netherlands; Department of Integrative Physiology, University of Colorado Boulder, Boulder, USA

**Keywords:** Daytime Light Exposure, Sleep, Circadian Phase, Slow Wave Activity, Sleepiness

## Abstract

**Study Objectives:** The causal effects of daylight exposure on sleep remain poorly understood, particularly under real-world conditions. This study aimed to address this gap by examining the effects of an afternoon daylight intervention on subsequent sleep.

**Methods:** Ten healthy participants (4 males; 19–28 years) completed two randomized 3-h afternoon outdoor walks under high-intensity light (HIL; clear goggles) or attenuated light (AL; neutral-density-filtered goggles; ∼10 photopic lux) conditions. Light exposure was centered around the estimated crossover between the phase-advancing and phase-delaying portions of the light phase-response curve, individually determined from baseline dim light melatonin onset (DLMO). Subsequent nocturnal sleep was assessed using at-home polysomnography (PSG), and subjective sleepiness and sleep quality were assessed using the Karolinska Sleepiness Scale and Groningen Sleep Quality Scale, respectively. Differences between conditions and across time were examined using linear mixed-effects models.

**Results:** There were no significant differences between conditions in DLMO timing or the phase angle between DLMO and sleep onset, indicating that the light manipulation did not produce a detectable shift in circadian phase. Sleep duration and self-reported sleep quality were also unchanged. However, nighttime delta power accumulated more rapidly following HIL exposure, accompanied by greater NREM sleep and reduced wakefulness after sleep onset (all p < 0.01). Participants were also significantly less sleepy upon awakening following HIL compared with AL exposure (p < 0.05).

**Conclusions:** High-intensity natural daylight exposure may promote greater sleep intensity and reduce sleepiness upon awakening. These findings highlight a potential role for daytime light exposure in modulating subsequent sleep physiology and next-morning sleepiness, independent of changes in circadian phase.

**Statement of significance:** Daytime light is typically thought to influence sleep primarily through its effects on timing of the circadian clock. Here, we show that three hours of high-intensity natural daylight altered subsequent sleep physiology without a detectable shift in circadian phase. High-intensity daylight was followed by deeper, less disrupted sleep and less sleepiness upon awakening the next morning. These findings suggest that daytime light can shape how the brain sleeps, even without shifting *when* we sleep, revealing a potentially distinct pathway through which daytime light influences sleep-wake regulation.

## Introduction

Light is the primary environmental signal synchronizing the human circadian system to the external light-dark cycle and plays an important role in the regulation of sleep and wakefulness (Dautovich et al., 2019). The effects of light on sleep are commonly attributed to its influence on the timing of the circadian clock. Depending on the biological timing of exposure, light can advance or delay circadian phase (Crowley et al., 2014; Khalsa et al., 2003; Kripke et al., 2007), which can subsequently alter the timing of sleep. Evening and early-night light exposure, for example, typically delays circadian phase and can shift sleep to a later time (Khalsa et al., 2003). However, whether light can influence sleep regulation independently of changes in circadian phase remains less well understood.

Sleep-wake regulation is classically described by the interaction of two processes: a circadian process (Process C) and a homeostatic process (Process S) (Borbély, 1982; Borbély et al., 2016). Process C reflects the approximately 24-hour variation in sleep-wake propensity generated by the circadian system, whereas Process S represents homeostatic sleep pressure, which progressively accumulates during wakefulness and dissipates during sleep. These processes can be characterized using complementary physiological and behavioral measures. Circadian phase can be assessed from markers such as melatonin or core body temperature (Borbély et al., 2016; Daan et al., 1984), while non-rapid eye movement (NREM) sleep EEG slow-wave activity (SWA) is considered a principal physiological marker of homeostatic sleep pressure (Borbély et al., 2016). During wakefulness, the progressive accumulation of homeostatic sleep pressure is also reflected in declining alertness and increasing sleepiness across time awake(Dijk et al., 1992; Lok, Woelders, Van Koningsveld, et al., 2022; Wright Jr et al., 2002).

Light could therefore influence sleep-wake regulation through more than one pathway. Its effects on circadian timing are well established: the magnitude and direction of light-induced circadian phase shifts depend strongly on the biological timing of exposure, as described by the phase-response curve (Crowley et al., 2014; Khalsa et al., 2003; Kripke et al., 2007). In contrast, considerably less is known about whether light exposure during wakefulness can influence the homeostatic regulation of subsequent sleep. If daytime light alters the accumulation of sleep pressure across wakefulness, this could be reflected in trajectories of alertness during the day and in the depth or intensity of subsequent sleep, even in the absence of a change in circadian phase.

Emerging evidence suggests that daytime light exposure may indeed be related to subsequent sleep. Outdoor light exposure is particularly relevant in this context because daytime outdoor illuminance can exceed 100,000 lux conditions, whereas typical indoor environments often provide less than 500 lux (Giménez et al., 2022). Observational studies have associated greater daytime light exposure with better self-reported sleep quality (Figueiro et al., 2017), greater time spent in NREM sleep (Wams et al., 2017), and longer actigraphy-derived sleep duration (Boubekri et al., 2014). Daytime light interventions have similarly been associated with improvements in subjective and objective measures of sleep (Ancoli-Israel et al., 2003; Viola et al., 2008). However, because light can simultaneously alter circadian phase, it remains unclear whether these associations reflect changes in circadian timing or whether daytime light can influence sleep regulation through mechanisms independent of circadian phase resetting.

One approach to disentangling these mechanisms is to administer light during a portion of the circadian phase-response curve in which phase-shifting responses are minimal. Around the crossover between phase delays and phase advances, light produces relatively little change in circadian phase (Crowley et al., 2014; Khalsa et al., 2003; Kripke et al., 2007). Exposure during this period therefore provides an opportunity to investigate the effects of daytime light on homeostatic sleep regulation while minimizing its influence on circadian timing.

The present study tested whether daytime light exposure influences sleep-wake regulation independently of changes in circadian phase. Daytime light exposure was experimentally manipulated in a real-world, home-based setting and timed to minimize circadian phase-shifting effects. Circadian phase and sleep timing were assessed to determine whether light altered the timing of the circadian and sleep-wake systems. In parallel, alertness was repeatedly assessed across the waking period to characterize changes associated with the accumulation of homeostatic sleep pressure, while subsequent sleep depth and NREM slow-wave activity were assessed as markers of homeostatic sleep regulation during sleep. Together, these measures allowed us to test whether daytime light exposure alters the build-up and subsequent dissipation of sleep pressure, even in the absence of a shift in circadian timing.

## Methods

### Participants

Participants were recruited through advertisements, flyers, and word of mouth. Ten healthy adults were included in the study.

Participants were excluded based on factors that could affect light sensitivity, sleep, or stability of timing of the circadian clock. Exclusion criteria included color vision deficiency (assessed using the Ishihara color vision test); use of photosensitizing medication, sleep medication, melatonin, or recreational drugs; smoking; high habitual caffeine consumption ( ≥5 cups/day); diabetes; and a body mass index (BMI) outside the range of 18–27 kg/m². Participants were also excluded if they reported shift work within the preceding 3 months, travel across ≥2 time zones within the preceding month, poor sleep quality (Pittsburgh Sleep Quality Index [PSQI] >5) (Buysse et al., 1989), or excessive daytime sleepiness (Epworth Sleepiness Scale [ESS] ≥18) (Johns, 1991). Additional exclusion criteria included wearing glasses and usage of oral contraceptive (women only).

### Medical Ethical Approval

All study procedures were approved by the Medical Ethical Research Committee of the University Medical Center Groningen, the Netherlands (NL62482.042.17), and were conducted in accordance with the Declaration of Helsinki. All participants provided written informed consent prior to participation and received financial compensation for their participation.

### Study design

The study used a randomized, counterbalanced, within-subjects design. Participants first completed two weeks of actigraphy to verify stable habitual sleep–wake timing, followed by an at-home dim light melatonin onset (DLMO) assessment (further described below) to determine baseline circadian phase. Participants were instructed to maintain a stable sleep– wake schedule throughout the experimental period.

Participants subsequently completed two experimental sessions separated by at least one week. During each session, participants completed a 3-hour afternoon walk while wearing either light-blocking ski goggles (further described below) or clear-lensed goggles. The order of the two conditions was counterbalanced across participants. Experimental sessions were scheduled on the same weekday and at the same time of day for each participant.

For each experimental session, participants started and ended their walk at the University of Groningen and were instructed to walk in the surrounding countryside. Participants could select their own route but were asked to follow approximately the same route during both experimental conditions. A ±15-minute margin around the scheduled return time was permitted. Participants wore an Actiwatch (Philips Respironics, Bend, OR, USA) on the non-dominant wrist throughout the walk to quantify activity levels. Route adherence and timing were monitored using GPS location data shared via WhatsApp (WhatsApp LLC, Menlo Park, CA, USA).

Subjective sleepiness was assessed hourly throughout the walk using an online version of the Karolinska Sleepiness Scale (KSS) (Åkerstedt & Gillberg, 1990). Participants who did not complete a scheduled assessment received a reminder by text message or telephone from a member of the research team.

Following the 3-hour walk, participants returned home wearing blue-light blocking glasses and completed an at-home DLMO assessment to determine circadian phase following each experimental condition. Saliva samples were collected every 30 minutes using pre-labelled collection tubes, with KSS ratings completed at each sampling time point.

During the DLMO assessment, a researcher of the study team visited the participant’s home to apply electrodes for overnight polysomnography (PSG). Participants were allowed to initiate and terminate sleep at their preferred times and were required to have no scheduled morning obligations the following day to allow unrestricted sleep. Upon awakening, participants completed the Groningen Sleep Quality Scale (GSQS) to assess subjective sleep quality (Leppämäki et al., 2003) and provided a final KSS rating.

## Measurements

### At-home dim light melatonin onset assessment

Circadian phase was assessed at home using dim light melatonin onset (DLMO). Participants remained at home under dim-light conditions approved by the study team. Curtains were closed, artificial light was restricted to a single low-intensity lamp, and participants wore blue-light-blocking glasses throughout the assessment. Light levels were additionally monitored using the light sensor of the MotionWatch 8 (CamNtech Ltd., Fenstanton, UK) to verify adherence to dim-light conditions.

The timing of salivary samples was individualized based on chronotype, assessed using the Munich Chronotype Questionnaire (MCTQ)(Roenneberg et al., 2003). Mid-sleep on free days corrected for sleep debt accumulated on workdays (MSFsc) was used to estimate habitual sleep timing. Dim-light conditions started 6 h before habitual sleep onset.

Saliva samples were collected every 30 min for a total of eight samples. Fifteen minutes before each scheduled sample, participants were instructed to remain seated and refrain from eating or drinking. Participants then provided a saliva sample by chewing on a cotton swab (Salivette; Sarstedt AG & Co. KG, Nümbrecht, Germany). While providing each sample, participants completed the KSS to assess subjective sleepiness. Pre-set timers were used to prompt sample collection. Following collection, samples were immediately refrigerated at approximately 4°C.

Upon awakening the following morning, participants notified the study team, and samples were transported to the University of Groningen for processing and storage (at −80°C until analysis). A baseline DLMO assessment was conducted before either experimental condition. Because circadian phase may differ between workdays and work-free days (Roenneberg & Merrow, 2016), baseline DLMO assessment and both experimental sessions were scheduled on the same day of the week for each participant.

### Salivary melatonin analysis

Saliva was collected using Salivette collection devices (Sarstedt AG & Co. KG, Nümbrecht, Germany). Salivary melatonin concentrations were quantified using a double-antibody radioimmunoassay (Direct Saliva Melatonin Radioimmunoassay; Bühlmann Laboratories AG, Schönenbuch, Switzerland). DLMO was defined as the interpolated time at which salivary melatonin concentration first exceeded 3 pg/mL. Linear interpolation between the last sample below and first sample above the 3-pg/mL threshold was used to estimate DLMO timing.

### Timing of light intervention

The timing of the light intervention was individualized relative to each participant’s circadian phase to minimize phase-shifting effects. The crossover region of the human phase-response curve, where the direction of the response to light transitions between phase delays and phase advances, was estimated to occur approximately 20 h before the core body temperature minimum (CBTmin) (Khalsa et al., 2003). CBTmin was estimated from DLMO using the previously described relationship for the Groningen student population: CBTmin = (DLMO − 20) × 1.25 + 3 (Woelders et al., 2017).

The 3-h light intervention was centered on the estimated crossover point, with the walk beginning 1.5 h before this time. Thus, light exposure was distributed approximately equally around the estimated transition between the phase-delay and phase-advance portions of the phase-response curve, with the aim of minimizing the net phase-shifting effect of the intervention.

### Light conditions

During both experimental conditions, participants wore clear-lensed ski goggles. In the attenuated-light (AL) condition, the goggles were fitted with multiple layers of neutral-density filters (LEE 211 0.9 ND 3 Stop Neutral Density; LEE Filters, Andover, UK) to reduce incident light intensity by approximately 3 log10 units. In the high-intensity-light (HIL) condition, participants wore otherwise identical goggles with clear lenses and without neutral-density filters.

### Actigraphy

Wrist activity was recorded in 1-min epochs using a MotionWatch 8 (CamNtech Ltd., Fenstanton, UK) worn on the non-dominant wrist. During the baseline period, actigraphy was used to verify regular sleep–wake timing; these recordings were not analyzed further. During experimental sessions, actigraphy quantified activity during the walking intervention, while the integrated light sensor monitored ambient light exposure. Recordings were analyzed using LightLoR (Zauner et al., 2025) The light sensor was also used to verify adherence to dim-light conditions during at-home dim-light melatonin onset (DLMO) assessments; these light recordings were not analyzed further.

### Home-based polysomnography

Sleep was assessed in-home using ambulatory polysomnography (PSG) with Actiwave recorders (CamNtech Ltd., Fenstanton, UK). The recording device was secured in a pouch worn by the participant to minimize interference with habitual sleep. The PSG montage included five scalp electroencephalography (EEG) electrodes positioned according to the International 10– 20 system at Fpz, C3, Cz, C4, and Oz; two electrooculography (EOG) electrodes; and one electromyography (EMG) electrode. The EMG electrode was positioned either beneath the chin or above the right eyebrow when facial hair or other factors prevented standard placement. A reference electrode was placed on the left mastoid. Electrodes were applied in participants’ homes by a researcher from the study team 2–3 h before habitual sleep onset. PSG signals were sampled at 128 Hz with 8-bit resolution.

### Polysomnographic processing and sleep scoring

PSG recordings were processed and visually scored using Embla RemLogic software (Natus Medical Incorporated, Middleton, WI, USA). Before scoring, signals were filtered using a 50-Hz notch filter, a 0.3-Hz high-pass filter, and a 35-Hz low-pass filter. Participants self-reported lights-off and lights-on times. Sleep stages were visually scored in 30-s epochs by an experienced sleep scorer who was blinded to participant identity and experimental condition. Sleep onset was defined as the first of three consecutive 30-s epochs scored as sleep (N1, N2, N3, or rapid eye movement [REM] sleep). Sleep outcomes included total sleep time (TST), wake after sleep onset (WASO), and time spent in N1, N2, N3, and REM sleep. Sleep-stage duration was expressed as a percentage of TST, due to the large variation in sleep duration among participants. WASO was used as an index of sleep disturbance and was expressed relative to TST.

### Polysomnography spectral analysis

PSG recordings from Fpz, Cz, and Oz were subjected to fast Fourier transformation (FFT) to quantify spectral power in the delta-frequency range (0.5–4.0 Hz) as a measure of slow-wave activity (SWA). Power spectral density was expressed in µV²/Hz. FFTs were calculated using 256-point windows with no overlap between consecutive windows. Epochs containing movement artifacts were identified and excluded from spectral analyses (Table S1). Across recordings, 20.4% ± 2.0% of NREM sleep epochs were excluded because of artifacts. SWA was quantified across NREM sleep for each electrode separately. To characterize the temporal dynamics of SWA across the sleep episode, spectral power was additionally averaged and accumulated across successive 1-h intervals and across 5% bins of total sleep time.

### Statistical analysis

Statistical analyses were performed in R [4.6.1; R Core Team, Vienna, Austria]. Linear mixed-effects models were fitted using the lme4 package to examine the effect of experimental light condition (HIL vs. AL) on subjective and objective sleep outcomes. Participant ID was included as a random intercept to account for repeated observations within participants. Outcomes included subjective sleep quality (GSS score), subjective sleepiness (KSS) upon awakening, sleep onset and offset timing, total sleep time, NREM and REM sleep duration, duration and percentage of TST spent in N1, N2, N3, and REM sleep, and NREM delta power and cumulative delta energy at Fpz, Cz, and Oz. Experimental light condition was included as the primary fixed effect.

To examine the trajectory of subjective sleepiness across the waking period, KSS ratings were analyzed using a separate linear mixed-effects model including experimental light condition, time, and their interaction as fixed effects, with participant ID included as a random intercept. Evening KSS measurements were additionally analyzed while accounting for the first evening KSS measurement.

Model selection and statistical significance were evaluated using Akaike’s Information Criterion (AIC) and Type II analysis of variance. Where appropriate, post hoc pairwise comparisons were performed using estimated marginal means with Tukey adjustment for multiple comparisons (emmeans package).

## Results

### General

The final sample consisted of 10 participants (6 females) with a mean age of 22.1 ± 2.5 years and a mean BMI of 22.60 ± 0.63 kg/m². Participants had a mean Pittsburgh Sleep Quality Index (PSQI) score of 3.70 ± 0.26 and a mean Epworth Sleepiness Scale (ESS) score of 7.60 ± 1.15. Mean chronotype, expressed as mid-sleep on free days corrected for sleep debt on workdays (MSFsc), was 4.38 ± 0.43 h. All female participants used oral contraceptives and completed both experimental conditions outside the hormone-free interval to minimize variation in hormonal status between conditions. Among the ten participants with complete paired light recordings, light exposure was significantly lower in AL than in HIL after applying the 3-log₁₀ attenuation correction to the wrist-recorded AL values (Table S2).

### Sleep Architecture

#### Cumulative delta power across sleep

There was no significant main effect of light condition on cumulative delta power in the frontal (F_(1,387)_ = 1.73, *p* = 0.18, Figure 2A), central (F_(1,387)_ = 2.22, *p* = 0.13, Figure 2B), or occipital locations (F_(1,387)_ = 2.20, *p* = 0.14, Figure 2D). As expected, cumulative delta power increased significantly across time asleep in all regions (frontal: F_(1,387)_ = 548.82, *p* < 0.001, Figure 2A; central: F_(1,387)_ = 477.73, *p* < 0.001, Figure 2B; occipital: F(_1,387)_ = 227.12, *p* <0.001, Figure 2C).

**Figure 1:**
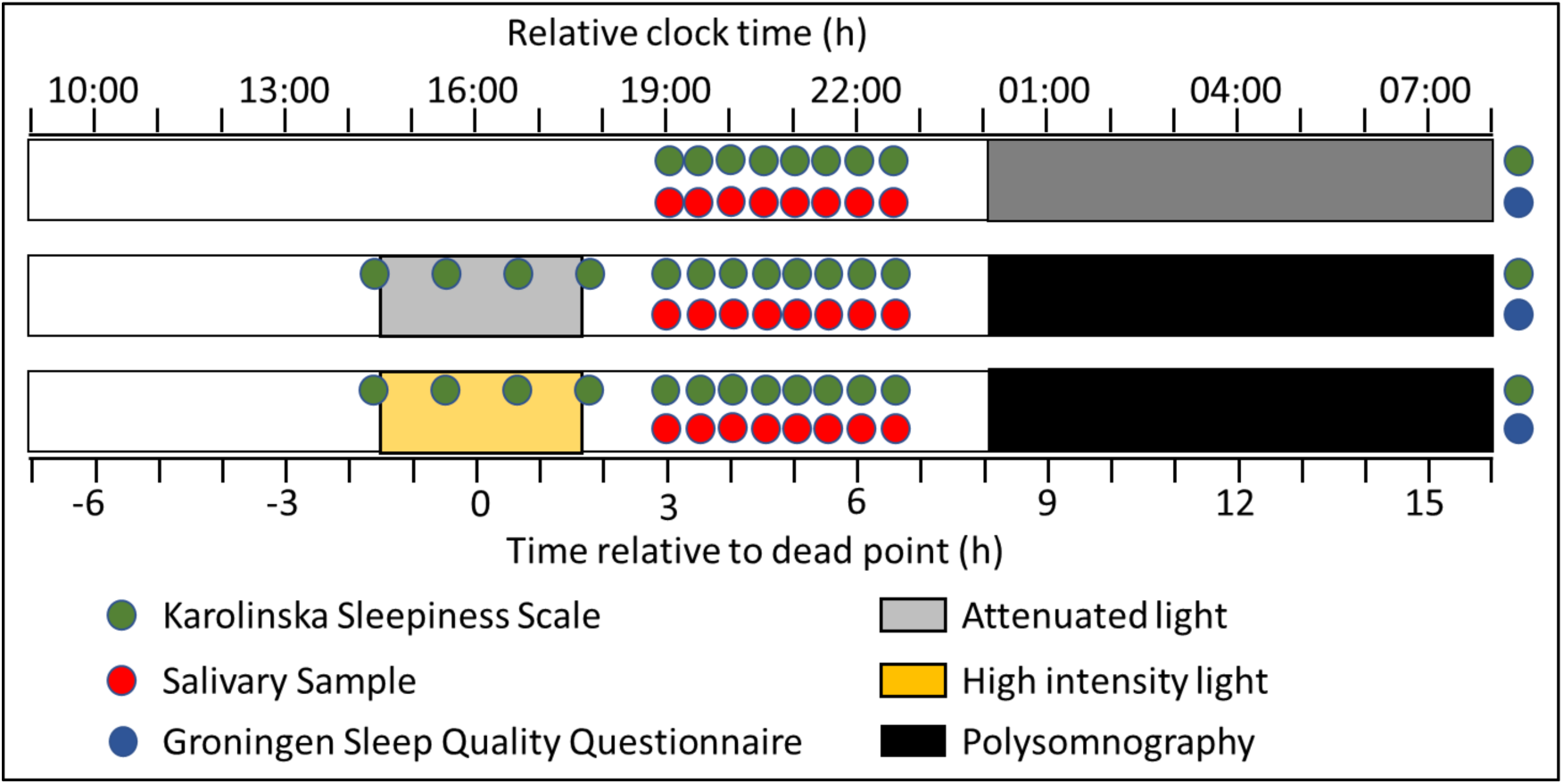
Experimental protocol timeline. The relative clock time is shown for a participant with a DLMO of 9 PM. The timing of sleep was self-selected during the experimental days. The study followed a counterbalanced within-subjects design; participants completed both conditions (attenuated and high-intensity light exposure) in randomized order.

**Figure 2.**
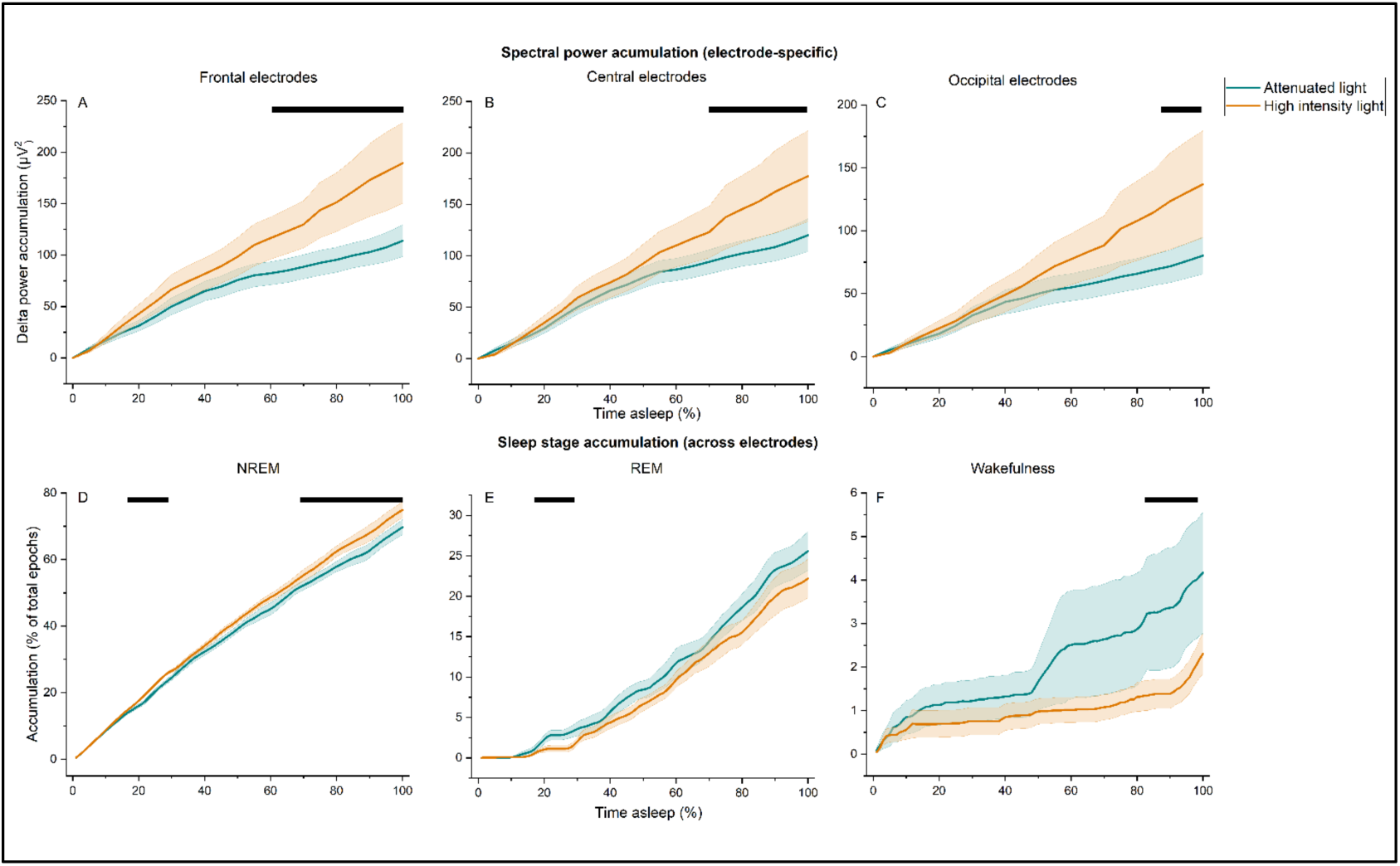
Spectral power and sleep stage accumulation. The upper panels depict the accumulation of delta power across experimental conditions in the frontal (A), central (B), and occipital (C) electrodes. The lower panels show the accumulation of sleep stages, including NREM sleep (D), REM sleep (E), and wakefulness (F). Data are presented separately for the attenuated light (teal) and high-intensity light (orange) conditions, with standard error of the mean presented in the same color. Significant differences between the two conditions are indicated by black bars.

Importantly, there was a significant time x light condition interaction for cumulative delta power in all regions (frontal: F_(1,387)_ = 42.95, *p* < 0.001, Figure 2A; central: F_(1,387)_ = 23.95, *p* < 0.001, Figure 2B; occipital electrode: F_(1,387)_ = 20.53, *p* < 0.001, Figure 2C), indicating that the accumulation of delta power across sleep differed between light conditions. Cumulative delta power increased more rapidly following HIL compared with AL exposure, with the difference between conditions becoming more pronounced later in the sleep episode (Figure 2A–C).

### Sleep Stage Accumulation

There was no significant main effect of light condition on cumulative NREM sleep (F_(1,1987)_ = 0.00, *p* = 0.99, Figure 2D), REM sleep (F_(1,1987)_ = 0.19, *p* = 0.66, Figure 2E), or wakefulness (F_(1,1987)_ = 0.51, *p* = 0.48, Figure 2F). A significant effect of time was observed for cumulative NREM sleep (F_(1,1987)_ = 65,322.48, *p* < 0.001, Figure 2D), REM sleep (F_(1,1987)_ = 9,102.97, *p* < 0.001, Figure 2E), and wakefulness (F(_1,1987)_ = 10,052, *p* <0.001, Figure 2F), indicating that sleep stage accumulation followed expected temporal patterns.

The temporal accumulation of sleep stages differed between light conditions. Significant time x light condition interactions were observed for NREM sleep (F_(1,1987)_ = 90.59, *p* < 0.001, Figure 2D), REM (F_(1,1987)_ = 31.99, *p* < 0.001, Figure 2E) and wakefulness (F_(1,1987)_ = 70.72, *p* < 0.001, Figure 2F). NREM accumulated more rapidly following HIL compared with AL exposure, with differences becoming more pronounced later in the sleep opportunity. Conversely, the accumulation of REM sleep and wakefulness was lower following HIL exposure, indicating a shift in the temporal distribution of sleep stages between the two light conditions.

Additional analyses presented in the Supplemental Information include individual hypnograms (Figure S1), the absolute time course of sleep (Figure S2), and comparisons of sleep-stage composition during the first 30, 60, 90, and 120 minutes of the sleep episode (Figure S3).

### Subjective sleepiness ratings Sleepiness during light exposure

Subjective sleepiness increased significantly across the 3-h walking period, as indicated by a significant main effect of time on KSS scores (F(_3,80_) = 5.15, *p* = 0.0032, Figure 3A). There was no significant main effect of light condition (F_(1,80)_ = 1.45, *p* = 0.23) and no time x light condition interaction (F(_3,80_) = 1.08, *p* = 0.364), indicating that the increase in sleepiness across the walking period did not differ between the HIL and AL conditions (Figure 3A).

**Figure 3.**
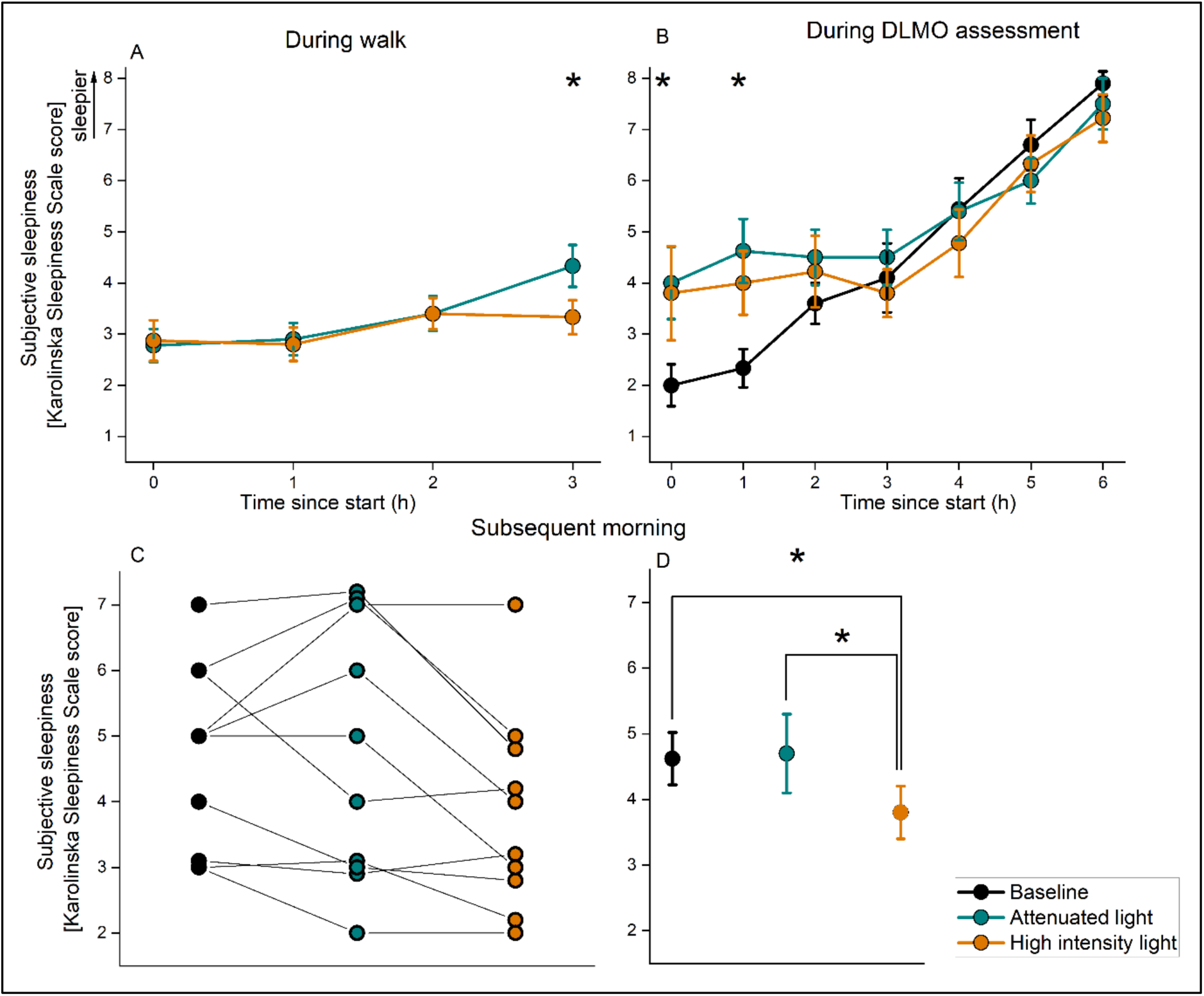
Subjective sleepiness across experimental conditions. (A) Karolinska Sleepiness Scale (KSS) scores during the 3-h light exposure, (B) during the subsequent dim light melatonin onset (DLMO) assessment, (C) and following awakening the next morning. Baseline, attenuated-light (AL), and high- intensity-light (HIL) conditions are shown in black, teal, and orange, respectively. Data are presented as mean ± SEM. Asterisks (*) indicate significant differences between conditions at individual time points.

### Sleepiness during DLMO assessment

KSS scores changed significantly across the DLMO assessment (main effect of time: F_(6,210)_ = 31.92, *p* < 0.001, Figure 3B). A significant main effect of condition was observed (F_(2,210)_ = 3.27, *p* = 0.041), as well as a significant time x light condition interaction (F_(12,210)_ = 2.04, *p* = 0.024), indicating that the temporal pattern of sleepiness differed between conditions. Post hoc comparisons using estimated marginal means indicated that sleepiness was higher at the first two measurement assessments following both experimental light conditions compared with the baseline assessment. By the end of the DLMO assessment, KSS scores did not significantly differ between conditions.

### Sleepiness the following morning

Morning KSS scores differed significantly between conditions (F_(2,30)_ = 3.69, *p* = 0.04, Figure 3C). KSS scores following the AL condition (4.7 ± 0.6) did not differ from baseline (4.63 ± 0.42; t = 0.05, p = 0.95). In contrast, KSS scores were significantly lower following the HIL condition (3.8 ± 0.4) compared with both baseline (t = −2.18, p = 0.044) and the AL condition (t = −2.44, p = 0.027), indicating lower subjective sleepiness the morning following HIL exposure.

### Circadian phase and timing of light exposure

At baseline, DLMO occurred at 21:08 ± 1:06 h (range: 19:13–22:53 h; Figure 4). The estimated core body temperature minimum (CBTmin) occurred at 04:25 ± 1:22 h (range: 02:02– 06:36 h), resulting in an estimated phase-response curve crossover point at 16:10 ± 1:22 h (range: 13:47–18:21 h). Accordingly, the 3-h light exposure began 1.5 h before this point, at 14:40 ± 1:22 h (range: 12:17–16:51 h), and ended at 17:40 ± 1:22 h (range: 15:17–19:51 h). DLMO occurred at 21:11 ± 1:39 h following the attenuated-light (AL) condition and at 20:58 ± 1:08 h following the high-intensity-light (HIL) condition (Figure 4). Relative to baseline, the mean phase shift was +3 ± 70 min following AL and −9 ± 42 min following HIL, corresponding to a mean difference of 12 ± 44 min between light conditions.

**Figure 4.**
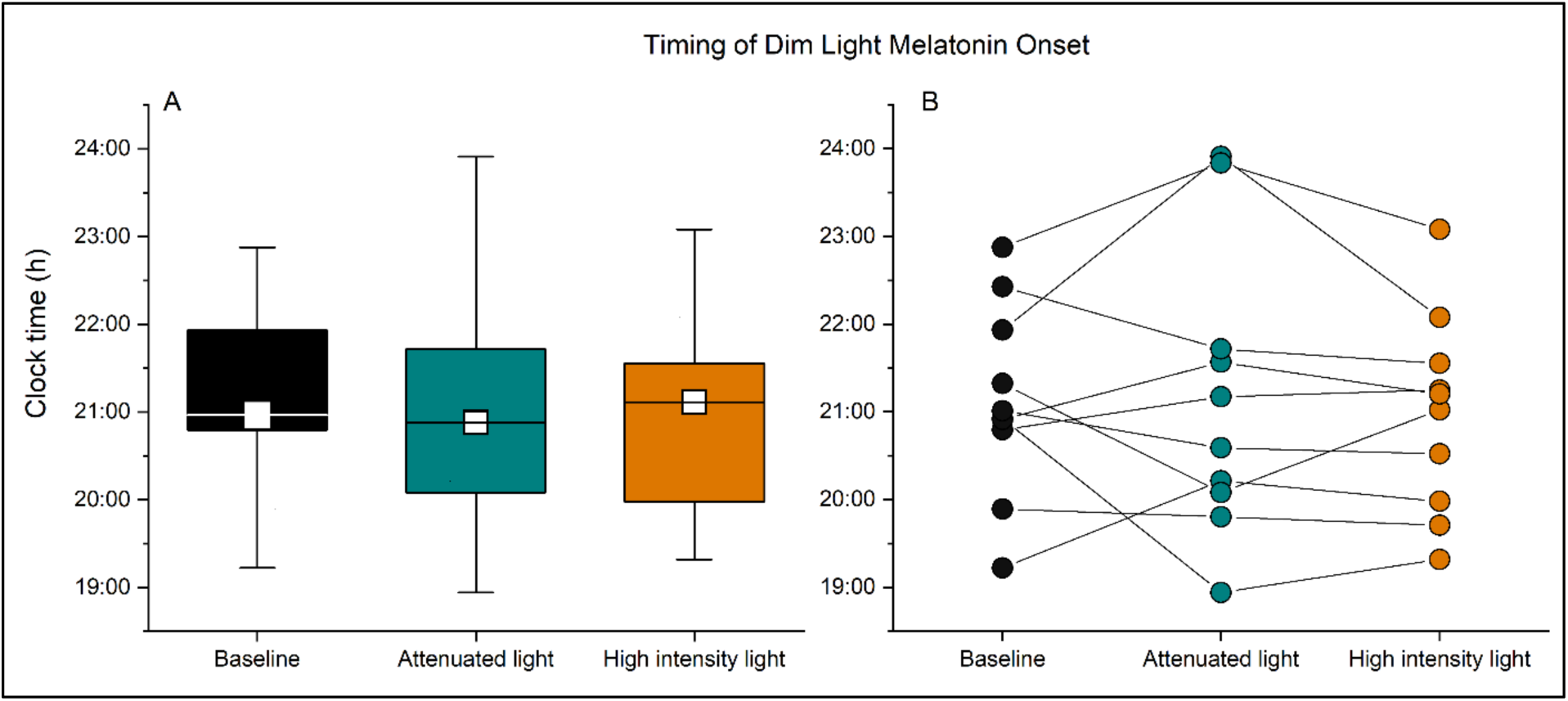
Timing of DLMO across experimental conditions. (A) DLMO timing at baseline (black), following attenuated light (teal), and high-intensity light (orange). (B) Individual changes in DLMO timing from baseline, following AL and HIL conditions. Individual participants are shown as points and lines

Light condition did not significantly affect DLMO timing (F_(2,30)_ = 0.30, *p* = 0.74; Figure 4), indicating that the experimental light manipulation did not result in a detectable shift in circadian phase.

### Self-reported sleep metrics

Self-reported sleep timing and duration did not differ between the HIL and AL conditions, including sleep onset (F_(1,20)_ = 1.94, *p* = 0.20, Figure 5A), offset (F_(1,20)_ = 1.16, *p* = 0.31, Figure 5B) or duration (F_(1,20)_ = 0.30, *p* = 0.59, Figure 5C). Self-reported sleep quality was also unaffected by light condition (F_(1,20)_ = 0.008, *p* = 0.93, Figure 5D). Similarly, the phase angle between DLMO and self-reported sleep onset did not differ between conditions (F_(1,20)_ = 1.68, *p* = 0.23, Figure 5E).

**Figure 5.**
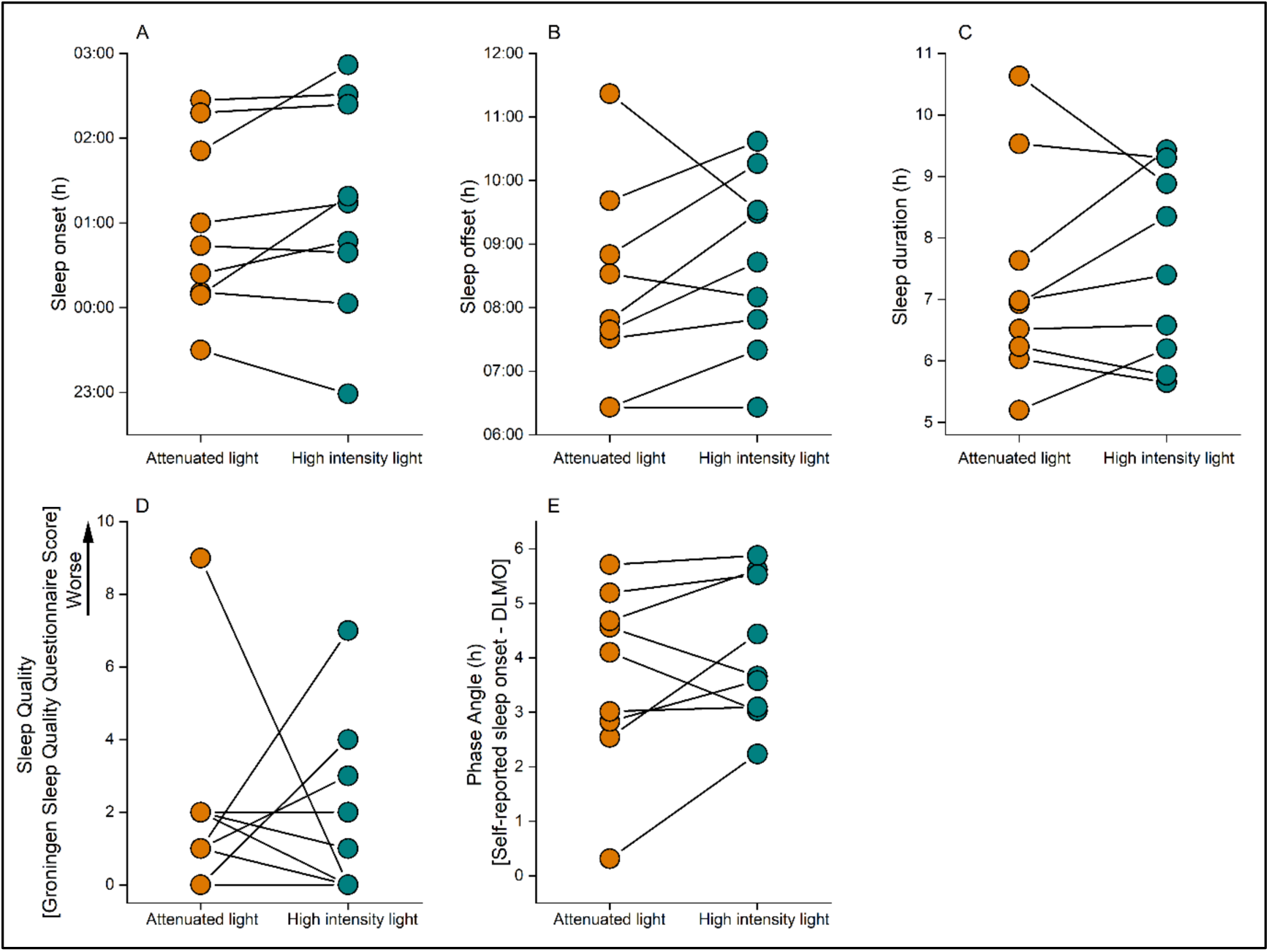
Self-reported sleep metrics across experimental conditions. Panels illustrate self-reported sleep timing and quality across experimental conditions. Presented are (A) self-reported sleep onset, (B) offset, (C) duration, (D) sleep quality, (E) and phase angle between DLMO and sleep onset. Attenuated light and high-intensity light conditions are presented in teal and orange, respectively.

## Discussion

The present study examined whether daytime light exposure can influence subsequent sleep and alertness without producing a detectable shift in circadian phase. Three hours of high-intensity natural daylight (HIL), timed around the crossover region of the phase-response curve, did not significantly alter DLMO timing, the phase angle between DLMO and sleep onset, self-reported sleep timing, sleep duration, or subjective sleep quality compared with attenuated light (AL). Despite this absence of detectable circadian phase shifting or changes in sleep timing, HIL exposure altered subsequent sleep physiology. Delta power accumulated more rapidly across sleep following HIL exposure, accompanied by greater NREM sleep accumulation and lower REM sleep and wakefulness accumulation. Participants also reported lower sleepiness the following morning after HIL exposure, without changes in self-reported sleep quality per se. Together, these findings suggest that daytime light may influence subsequent sleep-wake regulation through mechanisms that are not explained by circadian phase resetting alone.

A central feature of the experimental design was the timing of light exposure around the estimated crossover between the phase-advancing and phase-delaying portions of the human light phase-response curve(Khalsa et al., 2003). Consistent with the intended manipulation, DLMO timing did not significantly differ between the HIL and AL conditions (Figure 2). Similarly, neither the timing nor duration of subsequent sleep differed between conditions (Figure 4). Many effects of light on sleep are attributed to changes in circadian timing (Didikoglu et al., 2023; Dijk et al., 1999; Lok, Woelders, Van Koningsveld, et al., 2022). By experimentally minimizing phase shifting (Khalsa et al., 2003) and directly measuring DLMO, the present study provides preliminary evidence that daytime light can alter subsequent sleep physiology even in the absence of a detectable change in circadian phase.

Subjective sleepiness increased across the afternoon walk but, overall, did not differ between light conditions. This is consistent with previous studies reporting little or no effect of high-intensity light on subjective sleepiness at similar times of day (Borragán et al., 2017; Huiberts et al., 2016; Sahin et al., 2014). At the final time point, however, participants reported greater sleepiness following AL compared with HIL exposure, suggesting that an alerting effect of higher-intensity light may have emerged with prolonged exposure. As many previous studies have examined substantially shorter light exposures, often around 1 hour (Lok et al., 2018), it is possible that the effects of daytime light on subjective sleepiness develop cumulatively with increasing exposure duration. However, given that this difference was observed only at the final time point, this interpretation remains speculative and warrants further investigation.

At the beginning of the subsequent DLMO assessment, participants were sleepier following both light conditions compared with baseline, likely reflecting transient fatigue associated with the 3-h walk rather than an effect of light intensity (Merkelbach et al., 2011). This difference diminished across the evening as sleepiness in the baseline condition increased, with similar levels observed across conditions toward the end of the waking period. Thus, despite differences in the temporal trajectory of evening sleepiness, there was no clear evidence that HIL and AL differentially affected subjective sleepiness following the intervention.

The clearest physiological difference between conditions emerged during subsequent sleep. Although total sleep duration and subjective sleep quality were unchanged, the accumulation of delta power differed between conditions, with greater cumulative delta activity following HIL exposure. Delta activity during NREM sleep is closely associated with homeostatic sleep pressure and its dissipation across sleep. (Borbély et al., 2016). The accompanying differences in sleep-stage accumulation, with greater NREM sleep and less wakefulness following HIL, further indicate that daytime light exposure altered the organization of subsequent sleep (Lok, Woelders, Gordijn, et al., 2022). Importantly, these changes occurred without corresponding differences in DLMO or sleep timing. Although several studies have associated greater daytime light exposure with improved sleep (Wams et al., 2017), particularly in older adults (Bakker et al., 2004; Flores-Villa et al., 2020; Gimenez et al., 2014), these studies did not directly account for potential changes in circadian phase. The present findings therefore suggest that the effects of daytime light on subsequent sleep may extend beyond circadian phase shifting and involve processes related to sleep homeostasis or sleep intensity.

These effects on nocturnal sleep physiology were accompanied by a difference in subjective sleepiness the following morning. Participants were less sleepy upon awakening following HIL exposure than following both AL exposure and baseline, despite reporting no corresponding improvement in subjective sleep quality. While previous work has demonstrated that light administered around awakening can reduce sleep inertia (Gimenez et al., 2010; Santhi et al., 2013; Thompson et al., 2014; Van De Werken et al., 2010), the present findings suggest that daytime light exposure may have effects that persist across a subsequent night of sleep and influence sleepiness upon awakening. The absence of a perceived improvement in sleep quality despite changes in EEG-derived measures is not necessarily unexpected, as self-reported sleep quality does not always correspond closely with objective measures of sleep physiology (Kaplan et al., 2017; Lok, Chawra, et al., 2022; Lok et al., 2024).

The mechanisms underlying these effects remain unclear. An increase in homeostatic sleep pressure at sleep onset appears unlikely, as subjective sleepiness, which is related to homeostatic sleep-pressure accumulation (Czeisler & Gooley, 2007; Deboer, 2018), did not differ between light conditions toward the end of the evening, and subsequent sleep duration was unchanged. Nevertheless, the greater delta activity and NREM sleep, together with reduced wakefulness and REM sleep following HIL exposure, suggest that daytime light altered subsequent sleep intensity or organization.

Alternatively, light may have influenced aspects of circadian physiology other than phase. Experimental and modelling studies suggest that daytime light can increase circadian amplitude even when effects on circadian phase are minimal (Bano-Otalora et al., 2021; Walch et al., 2025), and greater daytime light exposure at these times of day has been associated with stronger SCN activity and reduced sleep-wake fragmentation (Bano-Otalora et al., 2021; Lok et al., 2023; Walch et al., 2025). However, circadian amplitude was not measured in this study, and this interpretation therefore remains speculative. Future studies combining measures of circadian phase and amplitude with markers of sleep homeostasis could determine the mechanisms through which daytime light alters subsequent sleep physiology.

Several limitations should be considered. First, the sample was relatively small (n = 10), and the findings therefore require replication in a larger and more diverse population. Second, the field-based design increased ecological validity but provided less control over environmental factors than a laboratory protocol. Although the within-subject design controlled for many effects associated with the outdoor walk itself, residual differences between conditions cannot be excluded. Third, the 3-log₁₀ attenuation correction was based on measurements at eye level, whereas ambient light exposure during the intervention was recorded at the wrist. Applying this correction to wrist-recorded illuminance therefore provides an estimate rather than a direct measurement of light exposure at the eye. Nevertheless, a 3- log₁₀ reduction represents a 1,000-fold decrease in illuminance (for example, from 10,000 to 10 lux) supporting a substantial exposure contrast between conditions despite uncertainty in the absolute eye-level values. Fourth, light exposure was specifically timed to minimize phase shifting; the findings therefore cannot be generalized to exposure at other circadian phases. Finally, although DLMO provided a direct measure of circadian phase, the study did not assess circadian amplitude or directly measure physiological markers of homeostatic sleep pressure during wakefulness.

## Conclusion

High-intensity daytime light exposure altered subsequent sleep physiology and reduced subjective sleepiness the following morning despite no detectable change in circadian phase, sleep timing, or sleep duration. Notably, these physiological changes were not accompanied by improvements in self-reported sleep quality, highlighting a dissociation between changes in measured sleep physiology and participants’ subjective perception of their sleep. Together, these findings provide preliminary evidence that the effects of daytime light on sleep-wake regulation extend beyond circadian phase resetting and may include modulation of processes involved in sleep homeostasis or sleep intensity. Larger controlled studies are needed to establish the underlying mechanisms and determine the functional relevance of these changes for next-day sleepiness and performance.

## Acknowledgments

MQY: data preprocessing and draft writing; TVB: data collection and draft writing; RL: study conceptualization, data analysis, and draft correction; RAH: study conceptualization, data analysis, and draft correction.

