## Supplemental Info for "Daylight exposure improves sleep and subsequent sleepiness"

**Supplementary information**

**Table S1. Overview of EEG artifacts across participants.** Across participants, 20.4% ± 2.0% of sleep epochs were identified as containing artifacts. In cases where EEG data from a specific channel were completely unavailable, data from an alternative channel were used for analyses. Specifically, FPz was replaced by Cz for S1 in the attenuated-light condition, Cz was replaced by C3 for S5 in the high intensity light condition and by C4 for S5 in the attenuated-light condition.

|  |  | Artifact Percentage (%) | | | | |
| --- | --- | --- | --- | --- | --- | --- |
| ID | condition | Cz | FPz | Oz | C3 | C4 |
| 1 | HIL | 9.17 | 9.17 | 9.17 | 9.17 | 9.17 |
| 1 | AL | 6.98 | 100.00 | 6.98 | 6.98 | 6.98 |
| 2 | HIL | 11.60 | 11.60 | 11.60 | 11.60 | 11.60 |
| 2 | AL | 10.65 | 10.65 | 10.65 | 100.00 | 100.00 |
| 3 | HIL | 21.18 | 12.32 | 12.32 | 12.32 | 31.90 |
| 3 | AL | 10.36 | 10.36 | 10.36 | 10.36 | 10.36 |
| 4 | HIL | 10.26 | 10.26 | 10.26 | 10.13 | 10.26 |
| 4 | AL | 12.46 | 12.46 | 12.46 | 12.46 | 100.00 |
| 5 | HIL | 100.00 | 6.61 | 6.61 | 6.61 | 100.00 |
| 5 | AL | 100.00 | 4.15 | 4.15 | 100.00 | 4.15 |
| 6 | HIL | 4.61 | 4.61 | 4.61 | 4.61 | 4.61 |
| 6 | AL | 5.65 | 5.65 | 5.65 | 5.65 | 5.65 |
| 7 | HIL | 5.89 | 5.89 | 5.89 | 5.89 | 5.89 |
| 7 | AL | 1.20 | 1.20 | 1.20 | 1.20 | 1.20 |
| 8 | HIL | 2.03 | 2.03 | 2.03 | 2.03 | 2.03 |
| 8 | AL | 6.77 | 6.77 | 6.77 | 6.77 | 6.77 |
| 9 | HIL | 24.19 | 24.19 | 24.19 | 24.19 | 24.19 |
| 9 | AL | 13.92 | 13.92 | 13.92 | 13.92 | 13.92 |
| 10 | HIL | 0.49 | 0.49 | 28.68 | 0.49 | 59.15 |
| 10 | AL | 0.36 | 0.36 | 0.36 | 0.36 | 30.92 |

**Table S2. Light exposure and activity metrics derived from MotionWatch8 recordings.** Metrics were calculated from one-minute recordings within each selected three-hour window. Light intensity in the attenuated-light condition (AL) was divided by 1,000 to account for the 3-log₁₀-unit attenuation; HIL values were unchanged. All light metrics were calculated using these corrected values. Cumulative light exposure represents light intensity integrated over the window, expressed in source light units × hours (lux·h if the source measurements are in lux). Median light intensity is the median of the one-minute readings. Light intensity SD describes overall variability within the window. The disparity index quantifies relative changes between consecutive readings, with higher values indicating greater minute-to-minute variability and zero indicating constant intensity. Total activity is the sum of activity readings in the device’s original units and was not adjusted for light attenuation. Time above threshold is the total number of minutes for which corrected light intensity exceeded the specified threshold. Paired comparisons included nine participants. Differences were calculated as HIL minus AL, with positive Cohen’s dᶻ values indicating higher values in HIL. Cohen’s dᶻ was calculated as the mean paired difference divided by the standard deviation of the paired differences. Two-sided Wilcoxon signed-rank and paired t-test p-values were adjusted using Holm’s method across the eight metrics, separately for each test.

| Metric | HIL mean | HIL SD | HIL median | AL mean | AL SD | AL median | Mean difference HIL - AL | Effect size (Cohen’s D) | Wilcoxon signed rank value (unadjusted) | Paired t.test (unadjusted) | Wilcoxen signed rank value (Holm-adjusted) | Paired t.test (Holm-adjusted) |
| --- | --- | --- | --- | --- | --- | --- | --- | --- | --- | --- | --- | --- |
| Cumulative light exposure | 25805.8 | 12445.2 | 25610.2 | 28.0858 | 9.09439 | 24.4989 | 25777.7 | 2.07109 | 0.00390 | 0.00025 | 0.03125 | 0.00102 |
| Median light intensity | 7091.68 | 3988.06 | 6407.12 | 7.67036 | 1.58051 | 7.83642 | 7084.01 | 1.77602 | 0.00390 | 0.00070 | 0.03125 | 0.00211 |
| Light intensity SD | 6719.80 | 2554.42 | 6739.79 | 7.78898 | 4.29765 | 6.81902 | 6712.01 | 2.62816 | 0.00390 | 4.85E-05 | 0.03125 | 0.00002 |
| Minute-to-minute light variability | 0.47056 | 0.10959 | 0.42304 | 0.35659 | 0.10818 | 0.36014 | 0.11396 | 0.64459 | 0.09765 | 0.08919 | 0.19531 | 0.17839 |
| Total activity | 73841.8 | 22281.6 | 70432.3 | 83579.6 | 25614.7 | 75848.4 | -9737.77 | -0.26296 | 0.65234 | 0.45291 | 0.65234 | 0.45291 |
| Time above 100 lux (min) | 174.888 | 10.6939 | 180 | 0.22222 | 0.66666 | 0 | 174.666 | 16.4494 | 0.00390 | 3.15E-11 | 0.03125 | 2.518E-10 |
| Time above 250 lux (min) | 173.444 | 11.3370 | 178 | 0 | 0 | 0 | 173.444 | 15.2989 | 0.00390 | 5.61E-11 | 0.03125 | 3.928E-09 |
| Time above 1,000 lux (min) | 164.111 | 17.9683 | 172 | 0 | 0 | 0 | 164.111 | 9.13335 | 0.00390 | 3.39E-09 | 0.03125 | 2.036E-08 |

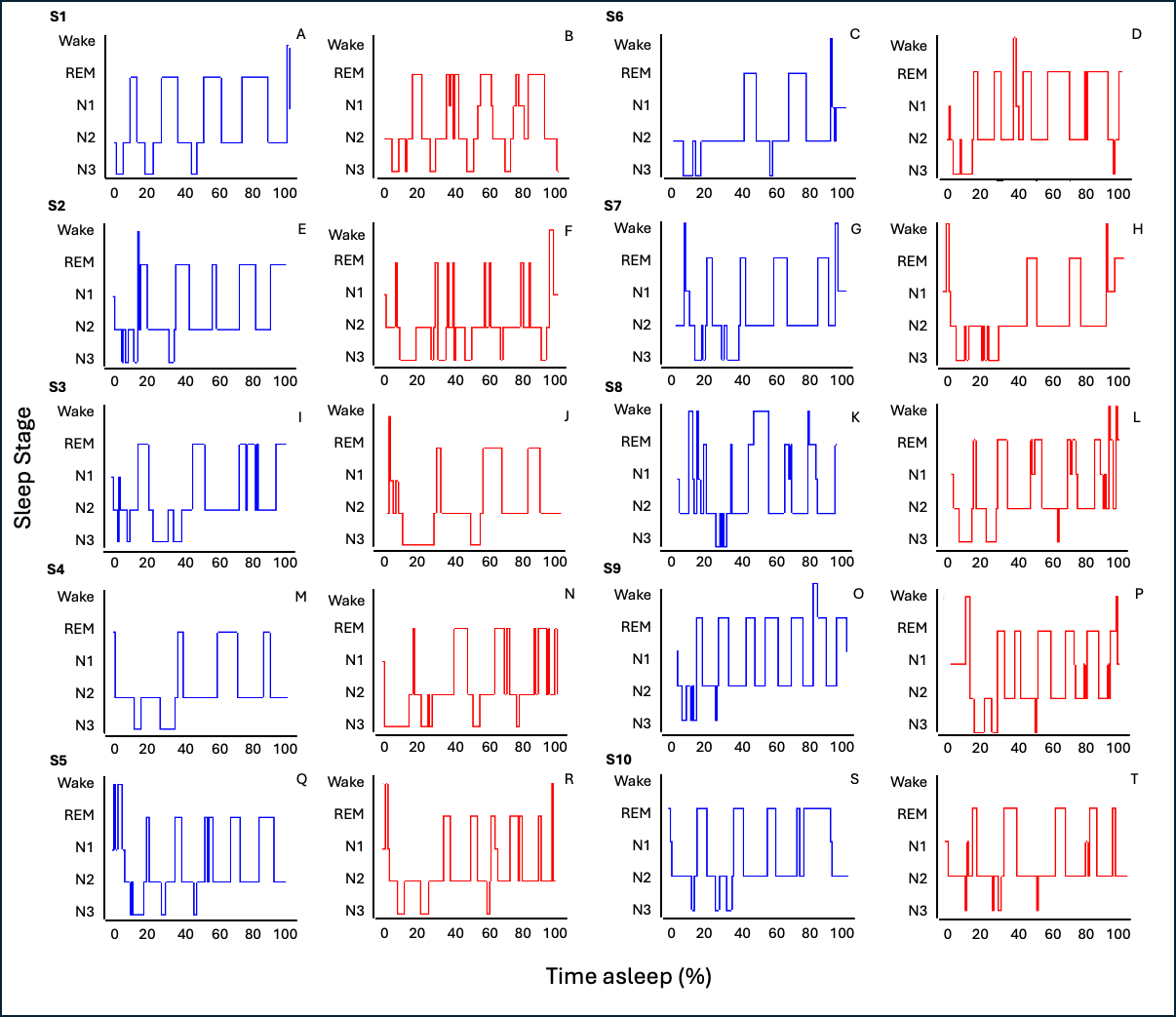

**Figure S1. Representative hypnograms illustrating sleep-stage transitions across the sleep period.** Hypnograms are shown for 10 participants (S1–S10), with two recordings displayed for each participant (blue, attenuated light condition; red, high intensity light condition). Sleep stages are plotted as a function of time asleep, normalized to the percentage of the total sleep period (0–100%). Sleep stages include wake, rapid eye movement (REM) sleep, and non-REM stages N1, N2, and N3.

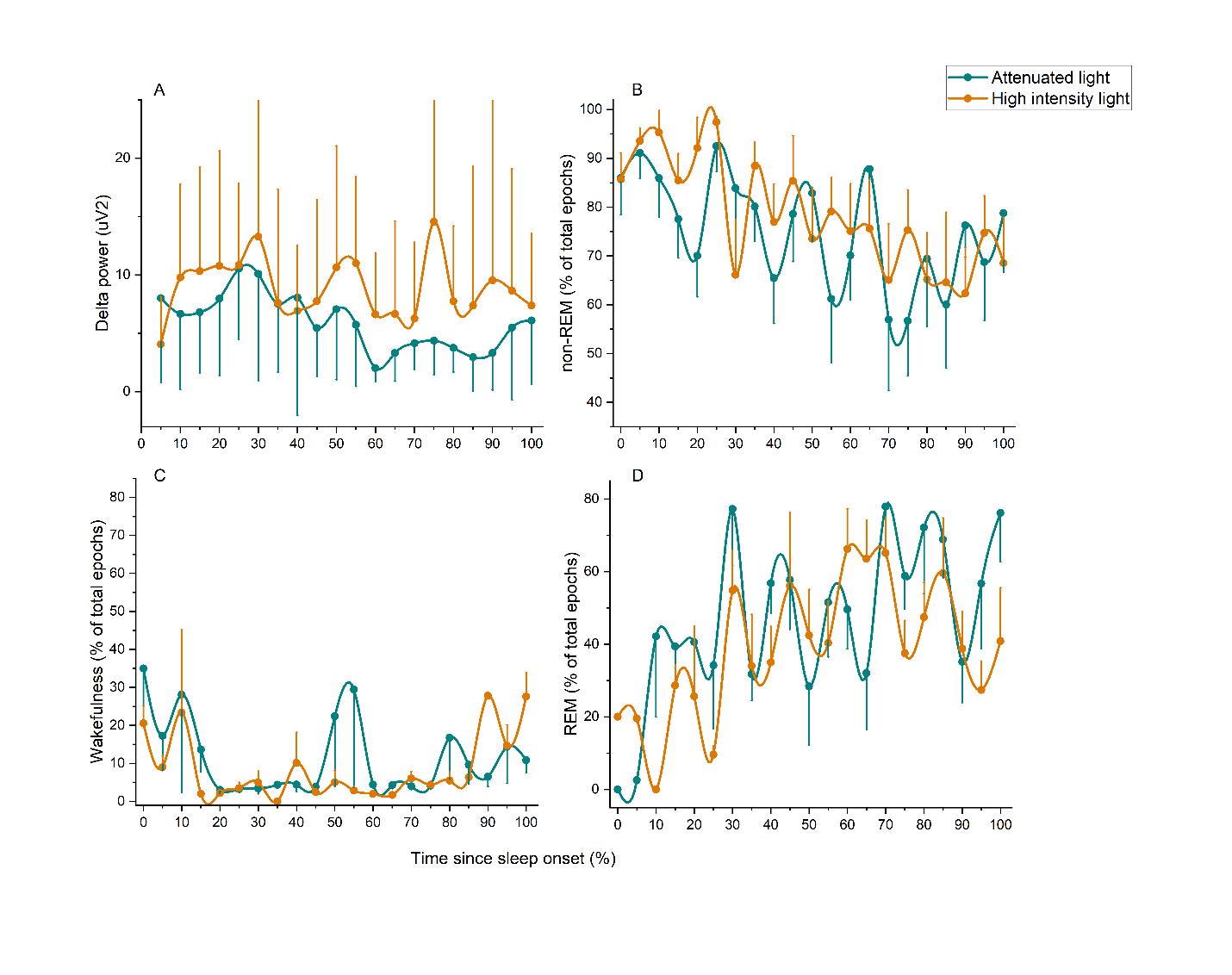

**Figure S2. Absolute time course of sleep parameters.** Time course of delta power (a), time spent in NREM sleep (b), REM sleep (C), and wakefulness (D) during the night, plotted relative to time since sleep onset. The attenuated light and high-intensity light conditions are presented in teal and orange, respectively. Error bars represent the standard error of the mean (SEM). Asterisks (*) indicate significant differences between conditions at specific time points.

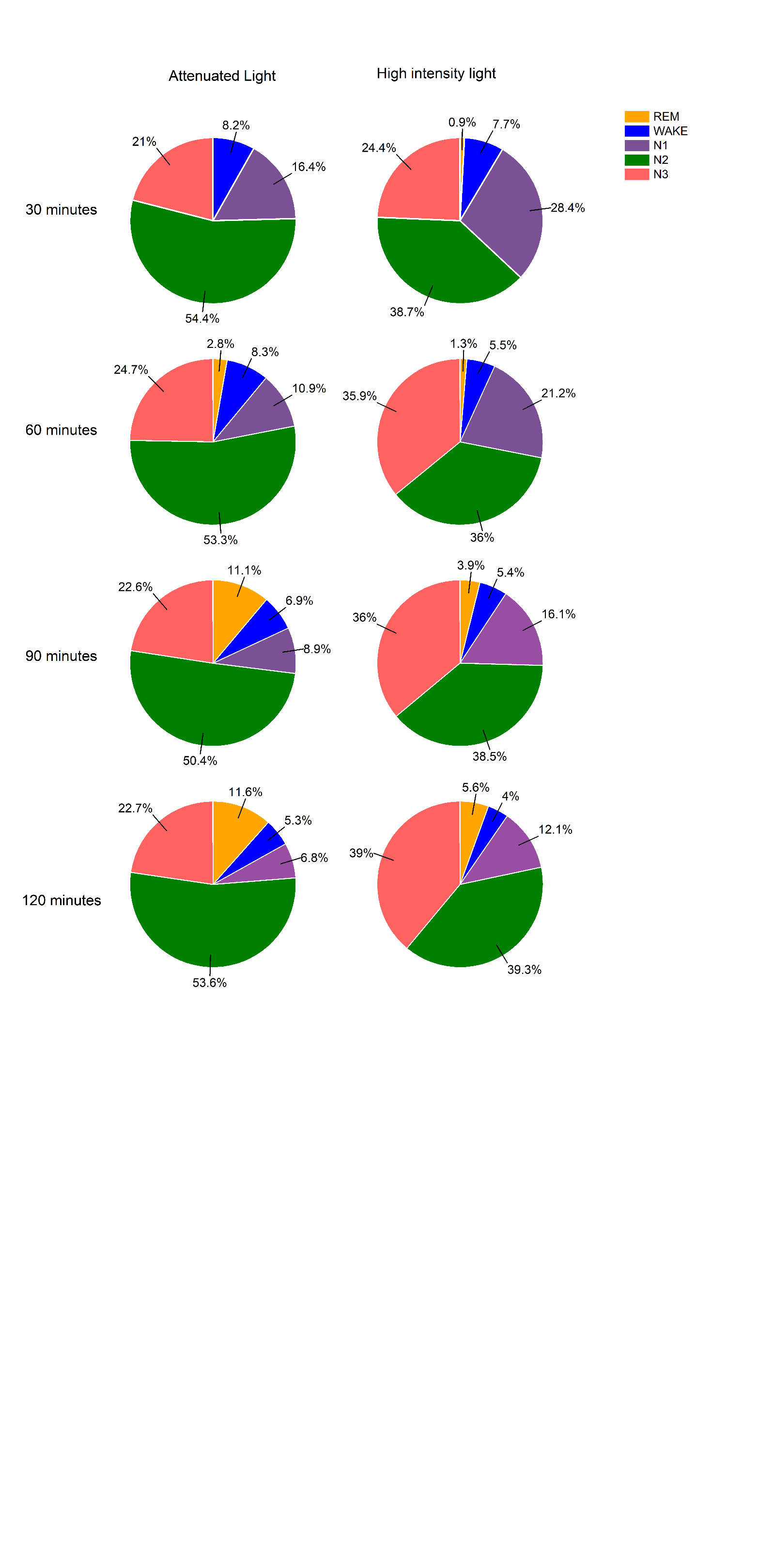

Figure S3. Sleep-stage composition under attenuated (left) and high-intensity (right) light conditions across increasing sleep durations. Pie charts show the percentage of time spent in each sleep stage during the first 30, 60, 90, and 120 minutes of sleep under attenuated light and high-intensity light conditions. Sleep stages are rapid eye movement (REM) sleep, wake, and non-REM sleep stages N1, N2, and N3. Percentages indicate the relative contribution of each sleep stage within the respective time interval. Sleep-stage composition was examined across progressively longer intervals to provide an indication of the dissipation of homeostatic sleep pressure over the course of the sleep period.
